# Bottlenecks can constrain and channel evolutionary paths

**DOI:** 10.1101/2022.07.15.500205

**Authors:** Jasmine Gamblin, Sylvain Gandon, François Blanquart, Amaury Lambert

## Abstract

Population bottlenecks are commonplace in experimental evolution, specifically in serial passaging experiments where microbial populations alternate between growth and dilution. Natural populations also experience such fluctuations caused by seasonality, resource limitation, or host-to-host transmission for pathogens. Yet, how unlimited growth with periodic bottlenecks influence the adaptation of populations is not fully understood. Here we study theoretically the effects of bottlenecks on the accessibility of evolutionary paths and on the rate of evolution. We model an asexual population evolving on a minimal fitness landscape consisting of two types of beneficial mutations with the empirically supported trade-off between mutation rate and fitness advantage, in the regime where multiple beneficial mutations may segregate simultaneously. In the limit of large population sizes and small mutation rates, we show the existence of a unique most likely evolutionary scenario, determined by the size of the wild-type population at the beginning and at the end of each cycle. These two key demographic parameters determine which adaptive paths may be taken by the evolving population by controlling the supply of mutants during growth and the loss of mutants at the bottleneck. We do not only show that bottlenecks act as a deterministic control of evolutionary paths but also that each possible evolutionary scenario can be forced to occur by tuning demographic parameters. This work unveils the effects of demography on adaptation of periodically bottlenecked populations and can guide the design of evolution experiments.

## 1 Introduction

Population bottlenecks are sudden, drastic reductions of population size that can arise both *in vivo* and *in vitro*. Pathogen populations experience such bottlenecks during host to host transmission [1, 2], or they can be induced by resource limitation or seasonality (e.g. the boom and bust dynamics of phytoplankton [3]). They also are commonplace in experimental evolution: in serial passaging (or transfer) experiments, a microbial population is periodically subsampled and placed on new medium to grow again [4]. This way, microbial populations can be followed during several generations [5] while remaining of a manageable size.

Several studies have investigated how periodic bottlenecks influence the rate of adaptation of such populations. They have mostly focused on the probability of stochastically losing beneficial mutations [6], on the time of arrival of successful mutations [7], on mutant fixation [8, 9, 10] and on the predictability of evolution [11, 12] with applications to the study of drug resistance [13, 14, 15] (see [16] for a review).

Here we examine theoretically how demography affects not only the rate of adaptation, but also the evolutionary paths followed by bottlenecked populations. In populations with constant size evolving in a regime of strong selection-strong mutation (as is often the case for experimental asexual populations), the distribution of fitness effect of fixed mutations and the rate of adaptation are dictated by the population size, the mutation rate and the shape of the distribution of fitness effects [17, 18]. In bottlenecked populations, both the supply of beneficial mutations and the probability of establishment change over time, and the accessibility of evolutionary paths can in theory depend on bottleneck size or intensity and on cycle duration. Furthermore, adaptation may increase population size, feed back on the mutation supply and enhance the scope for further adaptation. These effects are potentially important for experimental and natural evolution but have not been studied theoretically.

To study how bottlenecks influence the rate and paths of adaptation, we assume a minimal fitness landscape with a trade-off between the rate of appearance of beneficial mutations and their fitness advantage, consistently with the decreasing distribution of fitness effects documented in multiple species [19]. We take advantage of large population limit techniques as is standard in population genetics, notably in Luria-Delbrück type fluctuation experiments of microbial populations [20, 21]. We study the timing of emergence and establishment of beneficial mutations to investigate the effect of bottleneck size and cycle duration, or equivalently of the initial and final population sizes of the wild-type, on which paths are accessible to evolution.

## 2 Model

We consider an asexual population adapting to a new environment. We assume two types of beneficial mutations, with a trade-off between fitness and mutation rate. High-rate, weakly beneficial mutations confer a moderate gain in fitness, while low-rate, strongly beneficial mutations confer a large gain in fitness. This setting can be thought of as a coarse discretization of a decreasing (e.g. exponential) distribution of fitness effects [22]. These two types of mutations can be thought to target different loci underlying traits linked to adaptation, e.g. resistance to a drug or predator, or the exploitation of a new resource.

We thus obtain a simple fitness landscape composed of 4 genotypes: ‘00’ for wild-type (also denoted WT), ‘10’ for individuals with a weakly beneficial mutation, ‘01’ for individuals with a strongly beneficial mutation, and ‘11’ for individuals carrying both types of mutation. Thus, mutations 00 → 10 and 01 → 11 are weakly beneficial, while 00 → 01 and 10 → 11 are strongly beneficial. We allow magnitude epistasis but not sign epistasis, thus the growth rates of the four genotypes verify: *r*_11_ > *r*_01_ > *r*_10_ > *r*_00_. We neglect the production of double mutants by recombination between single mutants.

### A semi-deterministic model

Population size grows exponentially and is subject to periodic bottlenecks of fixed relative severity, i.e., the fraction of population that survives is constant (as opposed to fixed absolute bottleneck severity, where the number of individuals that survive is constant [16]). As a consequence, the WT population goes back to its initial size *N*_0_ at the beginning of each cycle. However, the size of the whole population can increase through successive transfers due to the arrival of beneficial mutants, see Figure 1. We use a semi-deterministic model to describe the dynamics of the population. We assume that *N*_0_ is sufficiently large that the growth of the WT population during one cycle can be described deterministically: *N_t_* = *N*_0_*e*^*r*_00_*t*^. In contrast, the population dynamics of mutants, always starting in small numbers of copies, will be described by a stochastic birth-death model.

**Figure 1:**
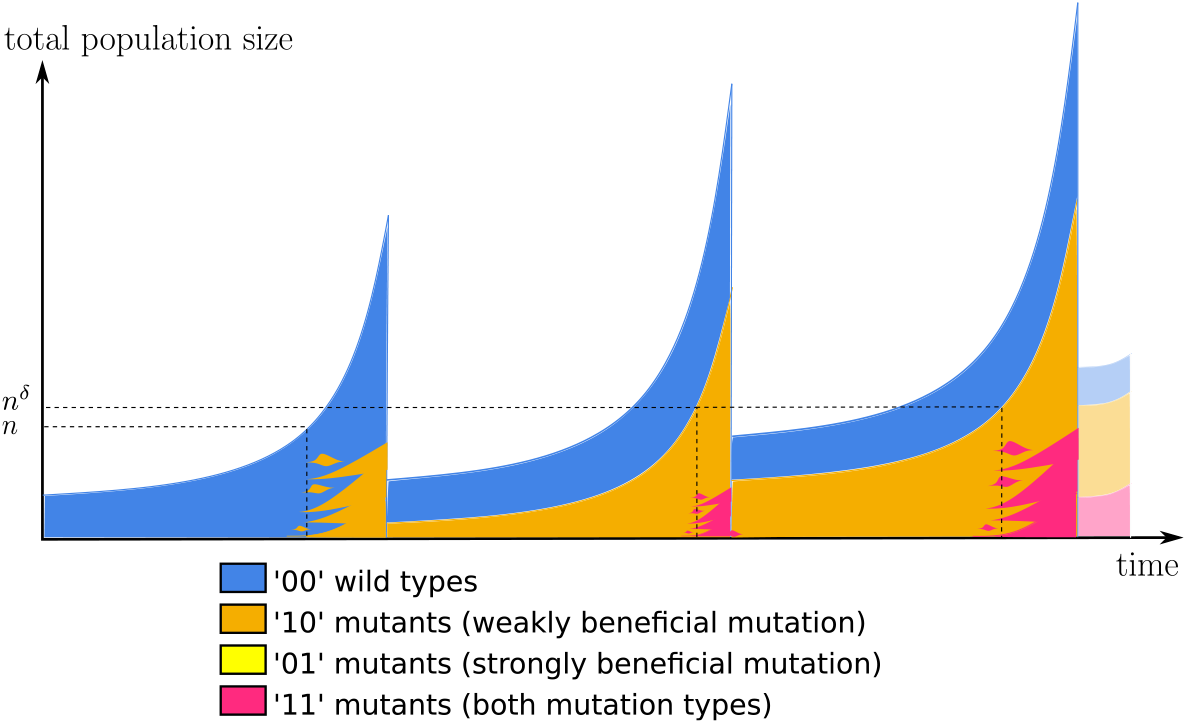
Illustration of a population evolving according to our model with periodic bottlenecks (3 cycles of growth are shown). Mutants start to appear when the population size is around the inverse of the mutation rate. Mutant subpopulations are composed of multiple independent clones, which can go extinct due to stochasticity (genetic drift, bottleneck). This scenario corresponds to the blue area in Figure 2, where strongly beneficial ‘01’ mutants never establish in the population but weakly beneficial ‘10’ mutants and double mutants do.

The weakly beneficial and strongly beneficial mutation rates are respectively *μ*_high_ and *μ*_low_, with *μ*_high_ ≫ *μ*_low_. We denote by *μ* = *μ*_high_ + *μ*_low_ the total mutation rate to beneficial mutations. This semi-deterministic setting is similar to the one first used in [23] to model the Luria-Delbrück experiment, and more recently in [24].

### Large population, small mutation rate assumption

The mutation rate to a beneficial mutation is typically very small, while the size of microbial populations is usually quite large. Thus we introduce a scaling parameter *n* that is of the order of 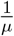, and we will assume in the following that *n* ≫ 1. In the large *n* limit the probability of most events of interest approaches either 1 or 0, allowing us to determine the most likely scenario in a given parameter setting.

To comply with *μ*_high_ ≫ *μ*_low_, we also introduce a parameter *δ* ∈ (1, 2) such that

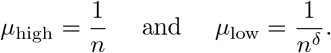

We suppose that the time *t_n_* between two dilutions is such that during one cycle of the experiment, the WT population grows from a size *N*_0_ = *n^β^* to *n^α^* with *β* ∈ (0, 1) and *α* > 1. As a consequence,

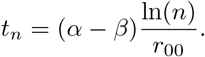

In this setting we have *N*_0_ ≪ *n* ≪ *N_t_n__*, ensuring that weakly beneficial mutants will appear during the first cycle with high probability. Because of the large *n* assumption, there is a sharp transition between a regime where mutations are very unlikely to occur (for *N_t_μ* ≪ 1) to a regime where numerous mutations arise (for *N_t_μ* ≫ 1). Thus we expect to observe adaptation via multiple-origins soft sweeps in the second regime [25, 26], in agreement with empirical observations in microbes [27, 28, 29].

The dilution factor between two cycles must be chosen so that the WT population always starts afresh at the same size *N*_0_ = *n^β^*. Thus the dilution factor is

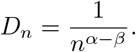

The notation is summarized in Figure 1 and Table 1.

**Table 1:** Main parameters of the model

| Notation | Interpretation |
| --- | --- |
| $r_{00}$ | growth rate of WT |
| $r_{10}, r_{01}$ | growth rates of single mutants |
| $r_{11}$ | growth rate of double mutant |
| $\mu_{\text{high}} = \frac{1}{n}$ | weakly beneficial mutation rate |
| $\mu_{\text{low}} = \frac{1}{n^\delta}$ | strongly beneficial mutation rate |
| $\mu = \mu_{\text{high}} + \mu_{\text{low}}$ | global beneficial mutation rate |
| $N_t$ | size of the WT population at time $t$ |
| $N_0 = n^\beta$ | initial WT population size |
| $N_{t_n} = n^\alpha$ | final WT population size |
| $t_n = (\alpha - \beta) \frac{\ln(n)}{r_{00}}$ | time duration of one cycle |
| $D_n = n^{-(\alpha - \beta)}$ | dilution factor |

## 3 Results

Here we analyze the dynamics of adaptation by characterizing the evolutionary paths followed by evolution, the timing of this process, and how it depends on demographic parameters. For detailed derivations, see Supplementary Information (SI), which also provides insight into the timing of weakly beneficial mutations during the first cycle.

When a mutant population arises at cycle *k*, escapes stochastic extinction and so is present in non-negligible quantity at the end of the growth phase, we will say that this population has **established** during cycle *k*. If the established mutant population reaches a sufficiently large size that it survives the next bottleneck (and so every subsequent bottleneck), we will say that this subpopulation **survives**. We say that an event A occurs with high probability (w.h.p.) if *P*(*A*) → 1 as *n* → ∞.

### 3.1 Effect of demography on evolutionary paths

#### Single mutant ‘10’ establishes w.h.p. at first cycle (assuming *α* > 1)

The product of the WT population size at the end of first cycle and of the high mutation rate is *μ*_high_*N_t_n__* = (1/*n*)*n^α^* ≫ 1. We rigorously showed in SI that ‘10’ mutants establish w.h.p. during the first cycle and computed an estimate of the number 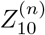 of ‘10’ mutants at the end of this cycle.

#### Single mutant ‘01’ establishes w.h.p. at first cycle if *α* > *δ* (and if *α* < *δ* w.h.p. never arises)

The establishment of mutants ‘01’ depends on the relative values of the final population size and the rate of strongly beneficial mutations, governed by parameters *α* and *δ*. If *α* > *δ*, then *μ*_low_*N_t_n__* ≫ 1 and mutants ‘01’ also establish during the first cycle. If on the contrary *α* < *δ*, the probability that mutants ‘01’ arise in the first cycle is close to 0. As the WT population has exactly the same size at the end of each cycle, it is unlikely that mutants ‘01’ establish in the course of the experiment. This highlights that the demographic control imposed on the WT population affects the establishment of mutations. In fact, it also affects which evolutionary paths are accessible: in the case where *α* < *δ* the transition 00 → 01 is not possible (and neither is 01 → 11).

#### Significance of demographic parameters *α* and *β*

For fixed values of the growth rates (*r*_00_, *r*_10_ and *r*_01_) and of *δ*, we can project on the plane (*β, α*) areas corresponding to different configurations of evolutionary paths. All path configurations must include the 00 → 10 transition because ‘10’ establish w.h.p. during the first cycle, thus there are 6 possible configurations. Figure 2 shows the areas corresponding to different path configurations for a chosen parameter set. This set was such that these 6 configurations are present, but this is not always the case. Equations for threshold lines (1-5) are derived and explained in SI.

**Figure 2:**
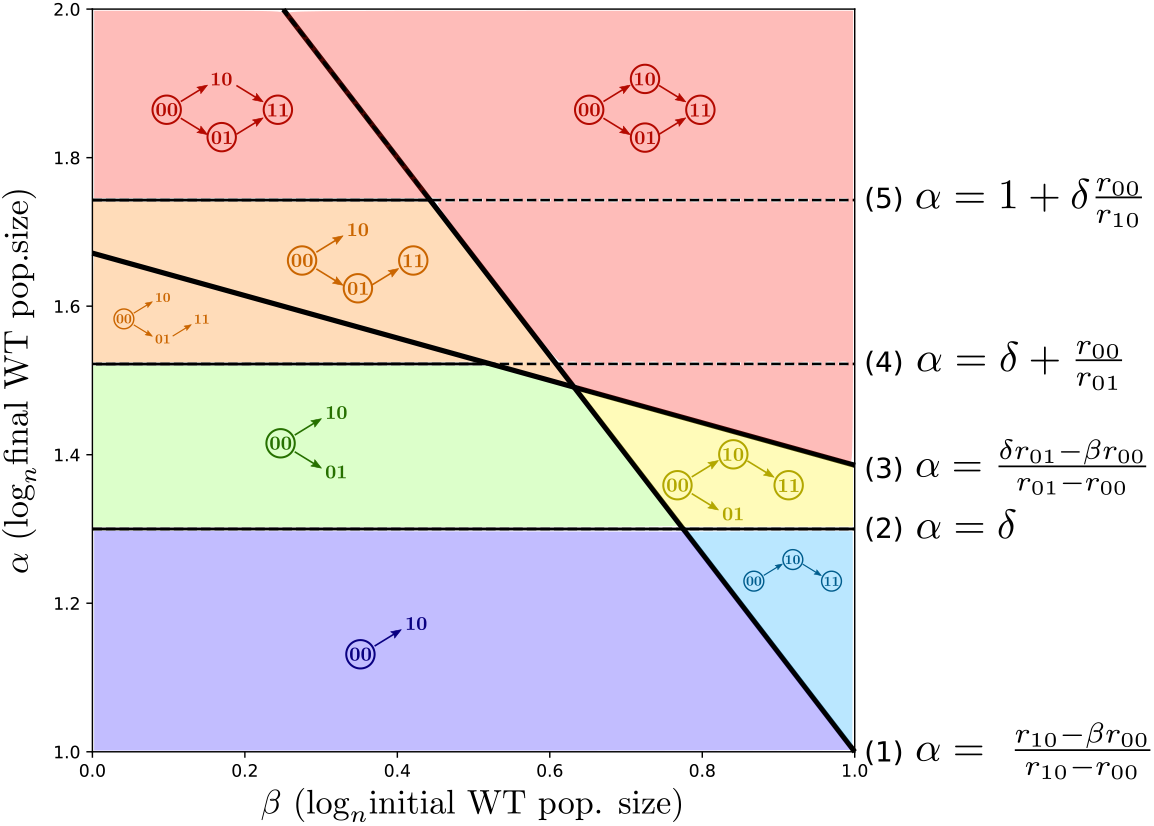
Predicted evolutionary paths, as a function of demographic parameters *β* and *α*. The six colors (red, orange, yellow, green, blue and purple) correspond to six different predicted path configurations. Genotypes that are shown but not circled are establishing but not surviving. Genotypes that are circled are surviving. The dilution ratio is constant along lines of slope 1. (*δ* = 1.3, *r*_00_ = 0.2, *r*_10_ = 0.35 and *r*_01_ = 0.9)

Together, *α* and *β* determine the initial and final WT population size at each cycle, the duration of each cycle and the relative severity of each bottleneck. When *α* increases, the final WT population size, bottleneck severity and cycle duration increase. When *β* increases, the initial WT population size increases but bottleneck severity and cycle duration decrease.

#### Single mutants survive if their population size after the first growth phase is larger than the inverse of dilution rate

Once a mutant population is established, it can either disappear because of the bottleneck or survive dilution and pass to the next cycle. For mutants ‘10’, these two outcomes correspond to two regions of parameter space delimited by line (1)

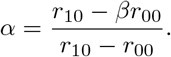

Indeed, *α* > (*r*_10_ – *βr*_00_)/(*r*_10_ – *r*_00_) is equivalent to having 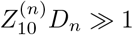 with 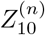 the number of ‘10’ mutants after one growth phase (see SI), and in that case the probability that at least one mutant ‘10’ survives dilution goes to 1 as *n* → ∞. Thus, above line (1) the ‘10’ mutant population survives the bottleneck w.h.p. and because the dilution factor is constant, it will also survive every subsequent bottleneck. The ‘10’ population will then continue growing, allowing double mutants to arise. Below this line, the ‘10’ mutant population at the end of the first cycle is too small to survive dilution, but re-establishes from the WT w.h.p. at each new cycle of the experiment.

For ‘01’ mutants the delimitation between establishment and survival is line (3)

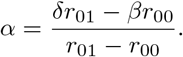

This line is analogous to line (1) except that mutant ‘01’ appears later (when population size reaches ~ *n^δ^*) but grows faster. Thus lines (1) and (3) delineate four different zones that we named the **Southwest, Southeast, Northwest** and **Northeast** corners. These zones can be further delimited into different colored areas depending on the value of *α* (Figure 2).

#### Southwest corner: no adaptation

In the Southwest corner, no single mutant population can survive. When *α* < *δ* (purple area under line (2)) only ‘10’ mutants can establish, they never survive but re-establish w.h.p. at each new cycle. Above line (2), in the green area, both ‘10’ and ‘01’ mutants establish at each cycle but w.h.p. never survive. When *α* increases above line (4), the time duration of a growth phase increases and the final population size of ‘01’ mutants becomes large enough for double mutants to arise and establish. These double mutants are lost in dilution w.h.p., except for high values of *r*_11_ for which they are able to survive even though neither ‘01’ nor ‘10’ survive themselves (case not represented on Figure 2).

#### Southeast corner: adaptation via ‘10’

In the Southeast corner, only mutants ‘10’ and ‘11’ survive. Depending on the sign of *α* – *δ*, we are either in the blue area with no ‘01’ mutant or in the yellow one with ‘01’ establishing repeatedly but not surviving bottlenecks. The example scenario displayed in Figure 1 corresponds to the blue area.

#### Northwest corner: adaptation via ‘01’

In the Northwest corner, only mutants ‘01’ and ‘11’ survive. The ‘10’ mutant population re-establishes at each cycle. If *α* increases above line (5), the growth phase lasts sufficiently long for the size reached by the ‘10’ mutant population to also produce double mutants.

#### Northeast corner: adaptation via both ‘10’ and ‘01’

In the Northeast corner (in red), all transitions are observed and all mutants will eventually establish and survive if enough cycles are performed.

### 3.2 Effect of demography on the timing of establishment of the double mutant

Here we show how demographic parameters affect the timing of adaptation, in particular at which cycle double mutants will establish. In Figure 2, lines (4) and (5) determine which value of *α* is needed for double mutants to establish during the first cycle. Above line (4) double mutants arise during the first cycle from mutant ‘01’, and above line (5) they arise during the first cycle from mutant ‘10’. Thus, as soon as *α* is above one of these two lines double mutants establish during the first cycle. Double mutants may also take more time to establish, as we now explain.

The different scenarios of double mutant establishment are illustrated in Figure 3, using the same parameter values as in Figure 2. Different shades of gray are used to indicate the speed at which the double mutants are predicted to establish. New lines (6-8) delineate these regions (see equations in SI).

**Figure 3:**
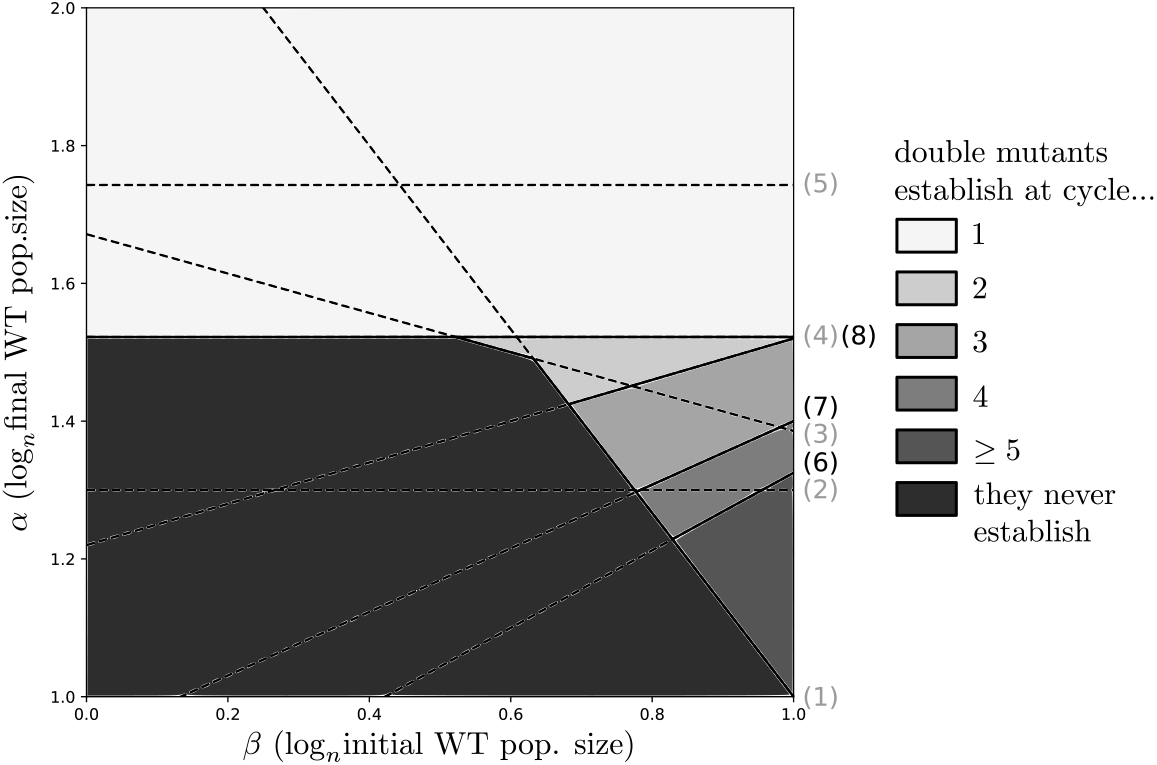
Predicted number of cycles to wait before the establishment of double mutants, as a function of demographic parameters *β* and *α* (*δ* = 1.3, *r*_00_ = 0.2, *r*_10_ = 0.35 and *r*_01_ = 0.9)

#### Black zone: double mutant never establishes

The black bottom-left zone located below lines (1), (3) and (4) corresponds to the green and purple areas of Figure 2 where single mutants re-establish w.h.p. at each cycle and cannot sustain a sufficiently large population to produce double mutants.

#### White zone: double mutant establishes at first cycle

In the white zone above line (4) double mutant establishes w.h.p. during the first cycle. It establishes either exclusively from mutant ‘01’ (between lines (4) and (5)), or from both ‘01’ and ‘10’ (above line (5)).

#### Grey zones: double mutant establishes after a finite number > 1 of cycles

In the remaining bottom-right zone, double mutants are predicted to establish in a number of cycles which is greater than 1 but w.h.p. finite. Lines (6), (7) and (8) indicate at which cycle single mutant populations are large enough for double mutants to establish.

### 3.3 Simulations with density-dependent division rate

In order to test our theoretical predictions in a more realistic setting, we performed simulations with density-dependent division rates: intrinsic division rates are multiplied by 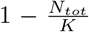, with *N_tot_* the current, total population size and *K* = *n^γ^* the carrying capacity. We place ourselves in the case *γ* > *α*: the WT population never reaches the stationary phase, but the mutant subpopulations may reach the stationary phase thanks to their higher growth rates (Figure S3 (SI)). In this setting, the dynamics are very similar to our first model without density-dependence, until one of the mutants reaches the stationary phase. Once the stationary phase is reached, in most cases the double mutant excludes competitively all other genotypes after a few cycles.

Figure 4 shows which scenarios were observed after simulating the evolution of a population with 1,000 different pairs of parameters (*α, β*) (details in SI). The color of a point corresponds to the evolutionary paths observed during the simulation, with the color coding used in Figure 2.

**Figure 4:**
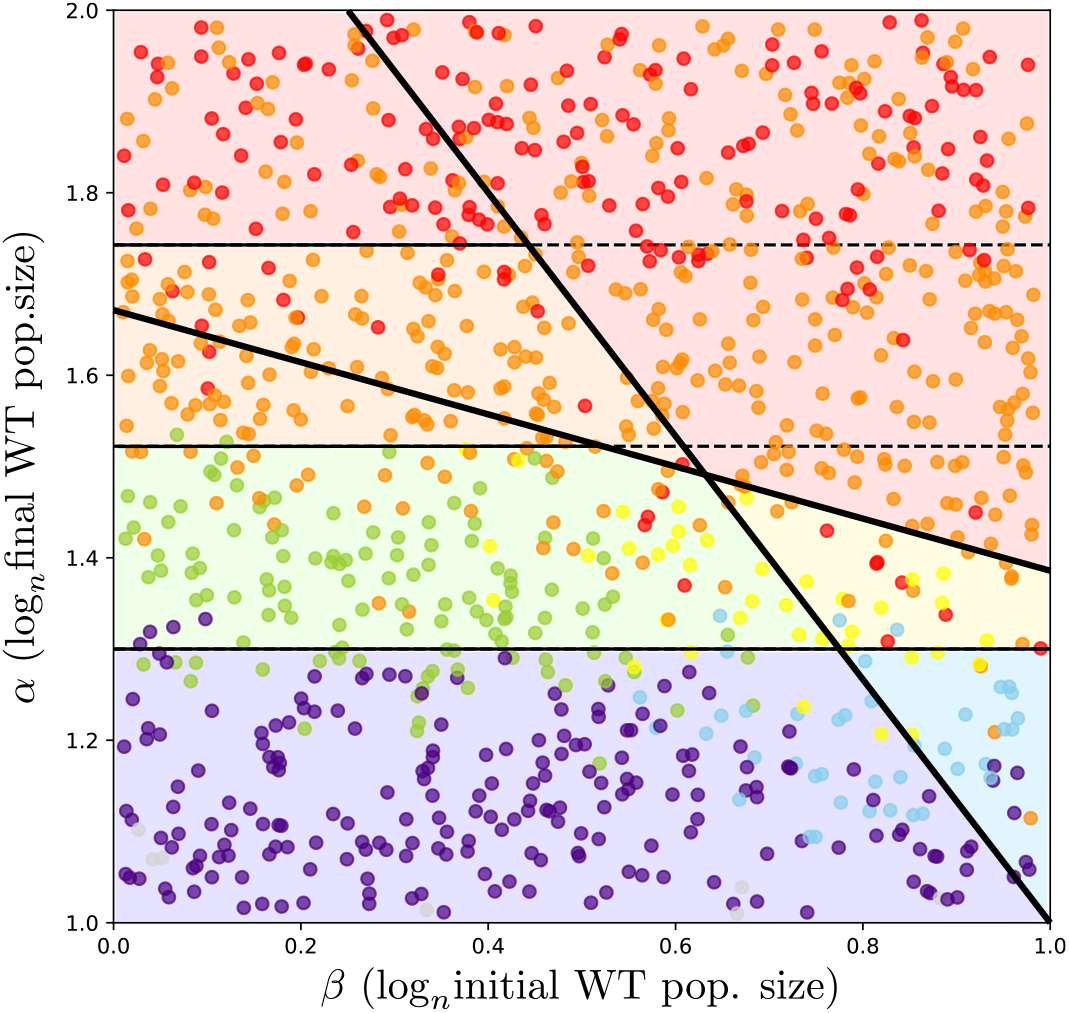
Observed paths in simulations with density-dependent division: each point corresponds to a simulation and is colored according to the observed scenario. Each simulation was run for 7 cycles, with parameters *n* = 10^12^, *γ* = 2, *δ* = 1.3, *r*_00_ = 0.2, *r*_10_ = 0.35, *r*_01_ = 0.9, *r*_11_ = 1 and death rates equal to 0.1. Boundaries and background colors are theoretical predictions for exponential growth and large *n* from Figure 2.

With density-dependence, it is still possible to observe the scenario where the establishment of the weakly beneficial mutation is followed by the establishment of the strongly beneficial mutation (blue area). Indeed, the establishment of the weakly beneficial mutation enables the population to reach the carrying capacity (*K* = *n^γ^*), allowing the emergence of the double mutant (as 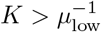).

However, the scenario where adaptation occurs through both weakly beneficial and strongly beneficial simple mutants (red area) is not observed because of competition: when both single mutants survive the first bottleneck, the weakly beneficial mutant ‘10’ is driven to extinction by the strongly beneficial mutant ‘01’ and thus is not able to produce double mutants. Although we still observe some red points on Figure 4, they correspond to situations where a small number of double mutants is produced from the weakly beneficial mutant at the first cycle, but do not survive dilution. Thus the population of surviving double mutants is only stemming from the strongly beneficial single mutant.

In conclusion, when mutants can reach a larger population size than WT, the only effect of density-dependence on evolutionary paths is to prevent the evolution of double mutants via two different paths.

## 4 Discussion

We characterize the trajectory and the speed of adaptation of an asexual, exponentially growing population subject to periodic bottlenecks. We studied adaptation on a minimal fitness landscape where only two classes of mutations are available: high-rate weakly beneficial mutations and low-rate strongly beneficial mutations.

Our main result is that 1) depending on initial and final population sizes, a unique evolutionary path unfolds and that 2) varying these two parameters, all paths can be explored. Establishment of a mutant is possible when the population size is of the order of the inverse of the relevant mutation rate. Surviving the bottleneck is possible when the final population size is of the order of the bottleneck severity. Tuning initial and final population sizes enables us to determine the evolutionary paths that the population will follow. A particularly interesting implication is that evolutionary paths can appear constrained not only because of sign epistasis [30] and rugged landscapes [31, 32, 33], but also because of fluctuating demography limiting the mutational input and causing the loss of beneficial mutations.

### 4.1 Effect of demographic parameters

Our model predicts which evolutionary paths will be observed, in the limit where the population size is large and the mutation rate is small. The predicted outcome depends both on the initial population size (β in logarithmic scale) and the final wild-type population size (α in logarithmic scale), where 1 in logarithmic scale corresponds to the inverse of the higher mutation rate. As *α* or β decreases, the accessibility of the double mutant decreases until it is no longer possible for the population to acquire both mutations. Indeed, decreasing population size at the end of the growth phase (decreasing *α*) limits the supply of mutations and increasing bottleneck severity (decreasing β) prevents mutant populations to survive until the next cycle.

Demographic parameters affect the rate of evolution. Increasing the final population size (α) always speeds up adaptation, as a large population size favors the emergence of mutations and also gives more time for the mutant subpopulation to reach a size large enough to survive the bottleneck. Interestingly, increasing the initial population size (β) has more complex effects. If the final population size is above the threshold for the establishment of the double mutant from the strongly beneficial ‘01’ mutant at the first cycle (α above line (4)), then the initial population size has no influence on the outcome. However, when the final population size is smaller than this threshold, establishment of the double mutant is fastest (in terms of number of cycles) for an intermediate initial population size. Indeed, when β is too small the bottleneck is too severe for mutations to survive. On the contrary if it is too large, then the bottleneck is less severe but the growth phase is shorter, leading to an overall effect of slowing down double mutant establishment. This non-monotonous effect of β is similar to that of the dilution ratio in [7]: a small ratio allows few mutations to survive, but a high ratio reduces the duration of a cycle and yields fewer mutations. However, increasing β decreases cycle duration hence the time to emergence of double mutant measured in absolute time (rather than in numbers of cycles).

### 4.2 Limitations

Our analysis relies on both a large population size and a small mutation rate approximation. However, these approximations correctly predict the outcome even for a finite population size. As shown on Figure S2, boundaries between different scenarios are blurred due to stochasticity but still visible. An important limitation of our model is that the population grows exponentially and is only bounded by the periodic bottlenecks, not by resource limitation. Coupled to the fact that the dilution ratio is kept constant throughout the experiment, this unlimited growth allows different clones to coexist indefinitely without interfering, which does not seem realistic [34, 35, 36]. If instead the total inoculum size were constant we would observe clonal interference [37, 38]. However, we do observe clonal interference when relaxing the assumption of exponential growth in a set of additional simulations with density-dependence. In this setting, the emergence of double mutants from both single mutants is no longer possible. Nevertheless, these simulations show that the rest of our results hold qualitatively when weakly beneficial mutants can reach a higher population size than wild-types. The mutants can reach a higher final population size than WT when the WT does not each stationary phase at the end of the cycle, as in our simulations. Alternatively, beneficial mutations can enable a larger stationary size: for example, in the Long Term Evolution Experiment the evolution of the ability to use citrate causes a ten-fold increase in final optical density, a proxy for population size [39].

### 4.3 Application to experimental design

Can our results be used to guide the design of evolution experiments? If mutation and division rates are known, it is possible to choose the size of the inoculum and bottleneck relative severity to decide which mutants emerge and when. Indeed, one can experimentally tune the values of α and β by first fixing the population size at the beginning of the experiment *N*_0_ = *n^β^* and then the duration of a cycle 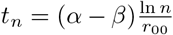. Recall that by assumption the dynamics of the WT population are periodic so that the dilution factor *D_n_* is fixed by the relation *D_n_* exp(*r*_00_*t_n_*) = 1. However all configurations may not be accessible in any given population or species, depending on the mutation rate and distribution of fitness effects. For example when *r*_01_ and *r*_01_ are close to *r*_00_, the lines (1) and (3) on Figure 2 are almost vertical. Thus the effect of periodic bottlenecks is most relevant when beneficial mutations confer a substantial fitness advantage, i.e., under strong selection.

In a more general setting where we have *k* beneficial mutations with a similar rate-benefit trade-off, and even without knowing precisely the mutation rate and distribution of fitness effects, what remains true is that increasing the initial and/or final population size will allow the population to access more diverse evolutionary paths. Furthermore, a large initial population size combined with a small final population size will favor frequent mutations, while a small initial population with a large final population size will favor rare but strongly beneficial mutations. The possibility to speed up the rate of adaptation by tuning demographic parameters could also alleviate the problem of bottlenecks in directed evolution [40, 41].

### 4.4 Interpretation of experiment outcome

Our results imply that sign epistasis is not necessary for evolution to follow a specific evolutionary path over others. For example, Figure 1 illustrates a scenario corresponding to parameter values falling into the blue area of Figure 2: the emergence of the strongly beneficial ‘01’ mutant is highly unlikely, but the weakly beneficial ‘10’ mutant establishes in the first cycle and the double mutant establishes during the second cycle. Without prior knowledge on traits and demography, an interpretation of the emergence of ‘11’ mutants exclusively from ‘10’ and never from ‘01’ mutants is sign epistasis: the fitness of the ‘01’ mutant is lower than the WT, but this mutation confers a benefit in the background of the other (weakly beneficial) mutation. However, this interpretation is incorrect here: the strongly beneficial mutation is beneficial in all backgrounds but emerges from ‘10’ and not from ‘00’, simply because ‘10’ mutants reach a higher population size than the WTs during the course of the experiment. The first set of beneficial mutations could thus enable access to other rarer mutations not through epistatic relationship but a larger final population size.

The phenomenon that we highlight here has been evidenced in experimental evolution. For example, Garoff et al. experimentally evolved *E. coli* under ciprofloxacin antibiotic in bottlenecks of varying severity [9]. They showed that for more severe bottlenecks, evolving mutations are weakly beneficial but affect mechanisms with large mutational target, for example efflux pump repressors. Rarer and more beneficial mutations only evolve when the bottleneck is less severe. A similar observation has been made by Schenk et al. for β-lactam antibiotic resistance [10]. This result is likely due to the bottlenecks preventing the emergence and/or survival of rare beneficial mutations.

All in all, these new mathematical results shed light on the factors shaping adaptation in repeatedly bottlenecked populations, showing that all paths can be followed by adaptation depending on demographic controls, and that the repeated appearance of specific evolutionary paths over others does not imply sign epistasis. This work calls for models studying the effect of demographic controls on evolution in more complex fitness landscapes and for inference methods disentangling the role of epistasis and demography in realized evolution experiments.

## 5 Acknowledgments

The authors are grateful to Guillaume Martin for several insightful comments. They thank the *Center for Interdisciplinary Research in Biology* (CIRB, Collège de France) for funding, as well as the MESRI (French Minister of Research) and the Ecole Polytechnique for the PhD scholarship of JG.

