## Supplementary Text for "Bottlenecks can constrain and channel evolutionary paths"

July 15, 2022

In this Supplementary Information we provide more details on our model (section 1), with proofs of results concerning the timing of mutations (sections 2 and 3) and the number of single mutants after one growth phase (sections 4 and 5). These results then allow us to derive the equations for threshold lines from Figure 2 and 3 of the main text (sections 6, 7, and 8). We also describe simulation results for exponential and density-dependent growths (section 9).

### Contents

**1 A semi-deterministic model** **2**
1.1 A few known results on birth-death processes . . . . . 2

**2 Waiting time to the first mutation** **3**

**3 Timing of weakly beneficial mutations** **3**

**4 Number of mutants '10' after one growth phase** **4**

**6 Survival of single mutants** **5**

**7 Constrained evolutionary paths** **6**

**8 Establishment of double mutants** **6**

**9 Simulations** **7**

### 1 A semi-deterministic model

As described in the main text, we use a semi-deterministic model in order to study the evolution of an asexual population experiencing periodic bottlenecks. The results presented in sections 2 to 5 of this Supplementary Information focus on the first growth phase of this population. During this phase, the wild-type population grows exponentially from a size $n^\beta$ ($\beta \in (0,1)$) at time 0 to a size $n^\alpha$ ($\alpha > 1$) at time $t_n$. The wild-type population size at time $t \in [0, t_n]$ is $N_t = N_0 e^{r_{00}t}$. $r_{00}$ is the growth rate and can be decomposed as $r_{00} = \lambda_{00}(1-\mu) - d_{00}$, with $\lambda_{00}$ the division rate, $\mu$ the beneficial mutation rate and $d_{00}$ the death rate.

We suppose that two categories of beneficial mutations can arise: weakly beneficial mutations with rate $\mu_{\text{high}} = \frac{1}{n}$ and strongly beneficial mutations with rate $\mu_{\text{low}} = \frac{1}{n^\delta}$ ($\delta > 1$). We focus here on weakly beneficial mutations and thus on mutants with genotype '10' (i.e. having a weakly beneficial mutation but no strongly beneficial mutation). Weakly beneficial mutations arise on the wild-type population according to a time-inhomogeneous Poisson process with parameter $\Lambda(t) = \lambda_{00}\mu_{\text{high}}N_t$. Each one of these mutations gives rise to a clone of mutants '10' following a birth-death process with birth rate $\lambda_{10}(1-\mu_{\text{low}})$ and death rate $d_{10}$.

#### 1.1 A few known results on birth-death processes

We denote by $Y_t$ the random variable measuring the size of a clone arising from a weakly beneficial mutation after a growth time of $t$. Then it can be shown (e.g. in [1]) that:

$$\mathbb{P}(Y_t = 0) = \frac{d_{10}(1-e^{-r_{10}t})}{\lambda_{10}(1-\mu_{\text{low}}) - d_{10}e^{-r_{10}t}}$$

$$\mathbb{P}(Y_t = k) = (1-\mathbb{P}(Y_t=0))\left(1 - \frac{\lambda_{10}(1-\mu_{\text{low}})}{d_{10}}\mathbb{P}(Y_t=0)\right)\left(\frac{\lambda_{10}(1-\mu_{\text{low}})}{d_{10}}\mathbb{P}(Y_t=0)\right)^{k-1} \quad \text{for } k > 0$$

We denote by $s_{10}(t)$ the probability for this clone to escape stochastic extinction at least until time $t$. Then:

$$s_{10}(t) = 1 - \mathbb{P}(Y_t = 0) = \frac{r_{10}}{\lambda_{10}(1-\mu_{\text{low}}) - d_{10}e^{-r_{10}t}}.$$

For $k > 0$, we can rewrite $\mathbb{P}(Y_t = k)$ as:

$$\mathbb{P}(Y_t = k) = (1-\mathbb{P}(Y_t=0))e^{-r_{10}t}s_{10}(t)\left(1 - e^{-r_{10}t}s_{10}(t)\right)^{k-1}.$$

Thus

$$\mathbb{P}\left(Y_t = k | Y_t > 0\right) = e^{-r_{10}t}s_{10}(t)\left(1 - e^{-r_{10}t}s_{10}(t)\right)^{k-1}.$$

This shows that $Y_t$ conditioned on survival is a geometric random variable with parameter

$$p_{10}(t) = e^{-r_{10}t}s_{10}(t) = \frac{e^{-r_{10}t}r_{10}}{\lambda_{10}(1-\mu_{\text{low}}) - d_{10}e^{-r_{10}t}}.$$

It follows that the expected clone size at time $t$ conditioned on the clone survival until $t$ is

$$\mathbb{E}\left(Y_t | Y_t > 0\right) = \frac{1}{p_{10}(t)}.$$

We denote this quantity as $c_{10}(t)$ in section 3.

#### 2 Waiting time to the first mutation

We denote by $T_n$ the time at which the first weakly beneficial mutation to escape stochastic extinction occurs. We define $\sigma_n$ as the time at which the wild-type population reaches a size of $n$ during the first growth phase. As the WT population size has an exponential growth $N_t = N_0 e^{r_{00}t}$ with $N_0 = n^\beta$, we have $\sigma_n = \frac{1-\beta}{r_{00}} \ln(n)$. In this section, we show that $T_n - \sigma_n = O(1)$ for $n \to \infty$. If we denote by $U_n$ the random variable equal to $T_n - \sigma_n$ and take $t > -\sigma_n$, then

$$\mathbb{P}(U_n \leq t) = \mathbb{P}\left(T_n \leq t + \sigma_n\right).$$

Weakly beneficial mutations arise according to a non-homogeneous Poisson process of parameter $\Lambda(t) = \lambda_{00}\mu_{\text{high}} n^\beta e^{r_{00}t}$, and the probability that a mutant '10' escapes stochastic extinction is $s_{10}(\infty) = \frac{r_{10}}{\lambda_{10}(1-\mu_{\text{low}})}$. Thus the probability that no such mutant occurs before time $t + \sigma_n$ is

$$\mathbb{P}\left(T_n > t + \sigma_n\right) = \exp\left(-\int_0^{t+\sigma_n} \Lambda(s) \frac{r_{10}}{\lambda_{10}(1-\mu_{\text{low}})}\,\mathrm{d}s\right).$$

Thus

$$\begin{aligned}
\mathbb{P}(U_n \leq t) &= 1 - \exp\left(-\int_0^{t+\frac{1-\beta}{r_{00}}\ln(n)} \lambda_{00}\mu_{\text{high}} n^\beta e^{r_{00}s} \frac{r_{10}}{\lambda_{10}(1-\mu_{\text{low}})}\,\mathrm{d}s\right) \\
&= 1 - \exp\left(\frac{\lambda_{00}\mu_{\text{high}} r_{10}}{\lambda_{10}(1-\mu_{\text{low}}) r_{00}} n^\beta \left(1 - e^{r_{00}t + \ln(n)(1-\beta)}\right)\right) \\
&= 1 - \exp\left(\frac{\lambda_{00}\mu_{\text{high}} r_{10}}{\lambda_{10}(1-\mu_{\text{low}}) r_{00}} n^\beta\right) \exp\left(-\frac{\lambda_{00}\mu_{\text{high}} r_{10}}{\lambda_{10}(1-\mu_{\text{low}}) r_{00}} n e^{r_{00}t}\right).
\end{aligned}$$

Recalling that $\mu_{\text{high}} = \frac{1}{n}$ and $\mu_{\text{low}} = \frac{1}{n^\delta}$ with $\delta > 1$, we obtain

$$\mathbb{P}(U_n \leq t) \underset{n\to\infty}{\to} 1 - \exp\left(-\frac{\lambda_{00} r_{10}}{\lambda_{10} r_{00}} e^{r_{00}t}\right).$$

The limit of the cumulative distribution function of $U_n$ shows that it converges in distribution to a random variable $U_\infty$, where $-U_\infty$ follows a Gumbel distribution with parameters $\left(\frac{1}{r_{00}} \ln\left(\frac{\lambda_{00} r_{10}}{\lambda_{10} r_{00}}\right), \frac{1}{r_{00}}\right)$. Thus we have:

$$T_n - \sigma_n \underset{n\to\infty}{=} O(1).$$

This shows that the first weakly mutation to escape stochastic extinction occurs around the time where the WT population size is of the order of the inverse of the mutation rate.

#### 3 Timing of weakly beneficial mutations

Here we look more closely at the times when weakly beneficial mutations occur during the first growth phase. We are interested in mutant clones that survive the first bottleneck, and we seek to investigate how their mutational origins are distributed over time. Mutational events giving birth to a surviving clone are called **surviving** mutations (see main text). The rate at which these mutations are produced at time $t$ during the first growth phase is

$$\lambda_{00}\mu_{\text{high}} N_t \propto e^{r_{00}t}.$$

Thus most mutations arrive at the end of the growth phase. Their distribution is plotted in pink on Figure 1. However, early mutations have more descendants. The final expected clone size (conditioned on survival) for a weakly beneficial mutation occurring at time $t$ is $c_{10}(t) = \frac{1}{r_{10}}\left(\lambda_{10}(1-\mu_{\text{low}})e^{r_{10}(t_n - t)} - d_{10}\right)$. The probability that a mutation occurring at time $t$ survives until the end of the growth phase being $s_{10}(t) = \frac{r_{10}}{\lambda_{10}(1-\mu_{\text{low}}) - d_{10}e^{-r_{10}(t_n-t)}}$, the rate of weakly beneficial mutations weighted by final expected clone size at time $t$ is

$$\lambda_{00}\mu_{\text{high}}N_t s_{10}(t)c_{10}(t) \propto e^{-(r_{10}-r_{00})t}$$

(see section 1 for a derivation of $c_{10}$ and $s_{10}$). This distribution is plotted in purple on Figure 1. The consequence is that clones that manage to survive the bottleneck have more chance to arise from a mutation arising in the middle of the growth phase (see solid blue curve on Figure 1). Indeed, if we suppose that the dilution is a binomial sampling, then the rate of surviving weakly beneficial mutations at time $t$ is:

$$\lambda_{00}\mu_{\text{high}}N_t s_{10}(t)\left(1 - \left(1 - n^{-(\alpha-\beta)}\right)^{c_{10}(t)}\right)$$

This distribution was already studied by Wahl et al. in a setting where the selective advantage is small [2]. They found that surviving mutations were likely to occur at all times during a growth phase, with a relatively flat distribution. This is indeed what we can observe on Figure 1 when $r_{10} \simeq r_{00}$ (dashed blue curve).

### 4 Number of mutants '10' after one growth phase

$Z_{10}^{(n)}$ is the random variable counting the number of mutants with genotype '10' at the end of the first growth phase. In the appendix, we show that $Z_{10}^{(n)} \underset{n\to+\infty}{\sim} n^{(\alpha-1)\frac{r_{10}}{r_{00}}} Z_{10}$. This shows in particular that mutants '10' **emerge** during the first growth phase, ie. they reach a large population size ($Z_{10}^{(n)} \gg 1$).

#### 4.1 Distribution of $Z_{10}$

The characteristic function of $Z_{10}$ is

$$\begin{aligned}
\phi_{Z_{10}}(t) &= \mathcal{L}_{Z_{10}}(-it) \\
&= \exp\left(-\frac{\lambda_{00}\pi}{\lambda_{10}\sin\left(\frac{r_{00}\pi}{r_{10}}\right)}\left(\frac{\lambda_{10}}{r_{10}}\right)^{\frac{r_{00}}{r_{10}}} |t|^{\frac{r_{00}}{r_{10}}} (-i\,\text{sgn}(t))^{\frac{r_{00}}{r_{10}}}\right) \\
&= \exp\left(-\frac{\lambda_{00}\pi}{\lambda_{10}\sin\left(\frac{r_{00}\pi}{r_{10}}\right)}\left(\frac{\lambda_{10}}{r_{10}}\right)^{\frac{r_{00}}{r_{10}}} \cos\left(\frac{\pi r_{00}}{2r_{10}}\right)|t|^{\frac{r_{00}}{r_{10}}}\left(1 - i\,\text{sgn}(t)\tan\left(\frac{\pi r_{00}}{2r_{10}}\right)\right)\right).
\end{aligned}$$

This expression tells us that $Z_{10}$ has a one-sided (Lévy) stable distribution supported by $[0,\infty)$ [3]. The stability parameter is $\frac{r_{00}}{r_{10}} \in (0,1)$, which implies that all moments are infinite. Sadly, the probability density function of $Z_{10}$ is not analytically expressible in the general case [4], nor are its median and mode.

### 5 Number of mutants ‘01’ after one growth phase

For mutants with genotype 01, there are two different cases depending on the sign of $\alpha - \delta$. Indeed, the probability that no mutant ‘01’ appears during a growth phase is:

$$\begin{aligned} p_{01}^{(n)} &= \exp\left(-\int_0^{t_n} \lambda_{00}\mu_{\text{low}} N_t \, \mathrm{d}t\right) \\ &= \exp\left(-\lambda_{00} n^{\beta-\delta} \int_0^{t_n} e^{r_{00}t} \, \mathrm{d}t\right) \\ &= \exp\left(-\lambda_{00} n^{\beta-\delta} \frac{1}{r_{00}} (n^{\alpha-\beta} - 1)\right) \\ &= \exp\left(-\frac{\lambda_{00}}{r_{00}} (n^{\alpha-\delta} - n^{\beta-\delta})\right) \end{aligned}$$

#### 5.1 Case $\alpha < \delta$

In this case, $p_{01}^{(n)}$ goes to 1 as $n \to \infty$. It is highly unlikely to see any ‘01’ mutant appear during the course of the experiment.

#### 5.2 Case $\alpha > \delta$

In that case, $p_{01}^{(n)}$ goes to 0 as $n \to \infty$. Furthermore, we can do the same computations as with mutants ‘10’ and find that the random variable $Z_{01}^{(n)}$ counting the number of mutants ‘01’ at the end of the first growth phase verifies:

$$Z_{01}^{(n)} \underset{n\to+\infty}{\sim} n^{(\alpha-\delta)\frac{r_{01}}{r_{00}}} Z_{01}$$

with $Z_{01}$ a stable distribution with stability parameter $\frac{r_{00}}{r_{01}}$. This shows that mutants ‘01’ emerge during the first growth phase.

### 6 Survival of single mutants

From Section 4 we know that mutants ‘10’ **establish** with high probability during the first cycle. However this does not guarantee that they will **survive**, that is, reach a sufficiently large size to survive the next bottleneck. Mutants ‘10’ survive if the quantity $A = -(\alpha-\beta) + (\alpha-1)\frac{r_{10}}{r_{00}}$ is positive. Indeed the expected quantity of mutants ‘10’ present at the beginning of the second cycle is $D_n Z_{10}^{(n)}$, with $D_n = n^{-(\alpha-\beta)}$ being the dilution factor. This quantity is equivalent to $n^{-(\alpha-\beta)+(\alpha-1)\frac{r_{10}}{r_{00}}} Z_{10}$ when $n$ is large, thus the population size of ‘10’ mutants starting in the second cycle is large if $A > 0$ and negligible if $A < 0$. If $A > 0$, the extinction probability of ‘10’ mutants is close to zero and their growth is deterministic for the remainder of the experiment, so that surviving one bottleneck entails the survival of all bottlenecks. If $A < 0$, with high probability no ‘10’ mutant survives the dilution as $n \to \infty$. Indeed, the probability that no mutant survives dilution knowing that $Z_{10} = z$ is

$$\left(1 - n^{-(\alpha-\beta)}\right)^{zn^{(\alpha-1)\frac{r_{10}}{r_{00}}}} \underset{n\to\infty}{\sim} \exp\left(-zn^{A}\right).$$

Then we discuss the **establishment** and **survival** of the strongly beneficial '01' mutant in the population. It establishes during the first cycle if $\alpha > \delta$. In this case, the number of '01' mutants at the end of the first cycle $Z_{01}^{(n)}$ is equivalent to $n^{(\alpha-\delta)\frac{r_{01}}{r_{00}}} Z_{01}$ for $n \to \infty$. Similarly to '10' mutants, they survive if $-(\alpha-\beta) + (\alpha-\delta)\frac{r_{01}}{r_{00}}$ is positive.
On the contrary if $\alpha < \delta$, then the probability that no '01' mutant is present at time $t_n$ $p_{01}^{(n)}$ goes to 1 as $n \to \infty$ (see section 5). As the WT population has the same growth trajectory in every phase, $p_{01}^{(n)}$ is in fact the probability to have no '01' mutant at the end of any cycle.

### 7 Constrained evolutionary paths

Parameters $\alpha$, $\beta$ and $r_{00}$ govern the demography of the wild-type population, while parameters $\delta$, $r_{10}$ and $r_{01}$ drive the appearance and growth rates of mutants. We have seen in section 6 that their relative values determine which mutations can survive, and a consequence is that they also determine which evolutionary paths are accessible. This is illustrated in Figure 2 of main text, where the different areas are delimited by the following threshold lines:

- (1) $\alpha > \frac{r_{10}-\beta r_{00}}{r_{10}-r_{00}}$ condition for the survival of mutants 10 (this is equivalent to $A > 0$);
- (2) $\alpha > \delta$ condition for the establishment of mutants '01' during first cycle;
- (3) $\alpha > \frac{\delta r_{01}-r_{00}}{r_{01}-r_{00}}$ condition for the survival of mutants 01;
- (4) $(\alpha-\delta)\frac{r_{01}}{r_{00}} > 1$ condition for the establishment of double mutants from mutant '01' during first cycle;
- (5) $(\alpha-1)\frac{r_{10}}{r_{00}} > \delta$ condition for the establishment of double mutants from mutant '10' during first cycle.

Conditions (4) and (5) are explained in the following section.

### 8 Establishment of double mutants

Demographic parameters constrain evolutionary paths, but also the timing of evolution. In particular, they determine in which cycle the double mutants '11' will establish. In the case where mutants '10' survive the first bottleneck, we can consider that their growth is deterministic during the following cycles. At the end of the $k^{th}$ cycle they have gone through $k-1$ dilutions and deterministic growth phases, thus their population size is

$$\begin{aligned} & n^{(\alpha-1)\frac{r_{10}}{r_{00}}} Z_{10} * D_n^{k-1} * (e^{r_{10}t_n})^{k-1} \\ & \underset{n\to\infty}{=} O\left(n^{(\alpha-1)\frac{r_{10}}{r_{00}}+(k-1)\left(\frac{r_{10}}{r_{00}}-1\right)(\alpha-\beta)}\right) \end{aligned}$$

Thus double mutants produced from mutant '10' establish during the first cycle $k_1$ such that

$$(\alpha-1)\frac{r_{10}}{r_{00}} + (k_1-1)\left(\frac{r_{10}}{r_{00}}-1\right)(\alpha-\beta) > \delta.$$

Similarly, double mutants produced from mutant '01' establish during the first cycle $k_2$ such that

$$(\alpha - \delta)\frac{r_{01}}{r_{00}} + (k_2 - 1)\left(\frac{r_{01}}{r_{00}} - 1\right)(\alpha - \beta) > 1.$$

As a result, double mutants establish at cycle $\min(k_1, k_2)$. This is illustrated in Figure 3 from main text, where additional lines are plotted compared to Figure 2:

- (6) $(\alpha - 1)\frac{r_{10}}{r_{00}} + \left(\frac{r_{10}}{r_{00}} - 1\right)(\alpha - \beta) > \delta$ condition for the establishment of double mutants from mutant '10' during the second cycle;
- (7) $(\alpha - 1)\frac{r_{10}}{r_{00}} + 2\left(\frac{r_{10}}{r_{00}} - 1\right)(\alpha - \beta) > \delta$ condition for the establishment of double mutants from mutant '10' during cycle 3;
- (8) $(\alpha - 1)\frac{r_{10}}{r_{00}} + 3\left(\frac{r_{10}}{r_{00}} - 1\right)(\alpha - \beta) > \delta$ condition for the establishment of double mutants from mutant '10' during cycle 4.

Lines representing conditions for the establishment of double mutants from mutant '01' during cycles 2, 3 and 4 are not plotted because they are below lines (6-8) and thus are not relevant for area delimitation.

### 9 Simulations

The code that was used to run the following simulations is available on Github at `https://github.com/JasmineGamblin/periodic-bottlenecks`.

#### 9.1 Exponential growth

We simulated populations evolving according to a fully stochastic model: individuals were grouped by subpopulations carrying the same genotype, each following a birth-death-mutation process. Double mutants originating from the weakly beneficial single mutant or the strongly beneficial single mutant were grouped separately. We used the principle of Gillespie's algorithm [5] to simulate the evolution of the system during growth phases, and binomial sampling to perform the dilution step. We resorted to a $\tau$-leaping approximation to speed-up the simulation when population sizes become large [6].

For each simulation, we recorded at the end of each growth phase which mutant subpopulations were present above a fixed threshold of 50 individuals (chosen so that the probability of extinction from this size is $< 1\%$). This way we could associate a color to the simulation according to the observed evolutionary paths, following the color coding from Figure 2 of main text.

We obtained Figures 2a)b)c) by randomly drawing 1,000 pairs of parameters $(\alpha, \beta)$ and simulating a population for 7 cycles, for different values of $n$. Populations for $n \geq 10^9$ and $\alpha > 1.4$ where simulated only for 4 cycles to avoid dealing with too large numbers. We can observe that for increasing values of $n$, the results of the simulations converge to the deterministic behavior that we predicted for $n \to \infty$. Grey points correspond to simulations where no mutant subpopulation was detected above the threshold.

##### 9.2 Density-dependent growth

For simulations with density-dependent division rates, we multiplied intrinsic division rates by $1-\frac{N_{tot}}{K}$, with $N_{tot}$ the total population size and $K=n^\gamma$ the carrying capacity. Remark that we choose to take the same carrying capacity for every subpopulation.

Figure 3 shows examples of simulated evolution for 11 different combinations of parameters $\alpha$ and $\beta$. We chose a parameter $\gamma=2$ such that the wild-types do not reach the stationary phase (because $\gamma>\alpha$). The total population size however can reach the stationary size due to the presence of beneficial mutants. Here the fact that growth is density-dependent introduces competition between the different subpopulations: we observe that wild-types and small-benefit mutants can be driven to extinction by fitter mutants.

Similarly to the exponential case, Figures 2d)e)f) were obtained by drawing 1,000 pairs of parameters $(\alpha,\beta)$ and simulating a population for 7 cycles, for different values of $n$.

Figure 4 was obtained by drawing 2,000 pairs of demographic parameters. For each pair, a population was simulated for 10 cycles and the cycle at which the double mutant emerges was recorded.

### A Appendix: Limit of $Z_{10}^{(n)}$ for $n\to+\infty$

Here we show that $Z_{10}^{(n)} \underset{n\to+\infty}{\sim} n^{(\alpha-1)\frac{r_{10}}{r_{00}}} Z_{10}$. We will first introduce some notations, then compute the Laplace transform of $\frac{Z_{10}^{(n)}}{n^q}$ and show that it converges to a non-trivial function for $q=(\alpha-1)\frac{r_{10}}{r_{00}}$ when $n\to\infty$.

#### A.1 Notations

We define the following random variables:

$M_{t_n}\in\mathbb{N}$ counts the number of weakly beneficial mutations arising among the WT population and still present at time $t_n$. $M_{t_n}$ is distributed as a Poisson random variable of parameter $\Lambda(t_n)=\int_0^{t_n}\lambda_{00}\mu_{\text{high}}N_t s_{10}(t_n-t)\,\mathrm{d}t$.

$T_1,\dots,T_{M_{t_n}}\in[0,t_n]$ are the times when these mutations arise. Conditioned on $\{M_{t_n}=m\}$, they are independent and identically distributed with probability density $\frac{\lambda_{00}\mu_{\text{high}}N_t s_{10}(t_n-t)}{\Lambda(t_n)}$.

$Y_t^{(i)}\in\mathbb{N}$ is the size after a growth time of $t$ of the clone arising from the $i^{th}$ weakly beneficial mutation. Conditioned on $\{M_{t_n}=m\}$, the $(Y_t^{(i)})$ with $i\in[1..M_{t_n}]$ are independent and identically distributed. $Y_t^{(i)}$ is a birth-death process starting at time $T_i$ and killed at time $t_n$, conditioned to survive until $t_n$. We recalled in section 1 that $Y_t^{(i)}$ is a geometric variable with parameter $p_{10}(t)=\frac{e^{-r_{10}t}r_{10}}{\lambda_{10}(1-\mu_{\text{low}})-d_{10}e^{-r_{10}t}}$.

With that we can express $Z_{10}^{(n)}$ as:

$$Z_{10}^{(n)}=\sum_{i=1}^{M_{t_n}} Y_{t_n-T_i}^{(i)}$$

#### A.2 Laplace transform of $Z_{10}^{(n)}/n^q$

Let us compute the Laplace transform of $\frac{Z_{10}^{(n)}}{n^q}$. For $u > 0$, we have:

$$\begin{aligned}
\mathcal{L}_{Z_{10}^{(n)}/n^q}(u) &= \mathbb{E}\left(e^{-u\frac{Z_{10}^{(n)}}{n^q}}\right) \\
&= \mathbb{E}\left(e^{-\frac{u}{n^q}\sum_{i=1}^{M_{t_n}} Y_{t_n-T_i}^{(i)}}\right) \\
&= \mathbb{E}_{M_{t_n}}\left(\mathbb{E}_{T_i|M_{t_n}}\left(\mathbb{E}_{Y_{t_n-T_i}^{(i)}|T_i}\left(e^{-\frac{u}{n^q}Y_{t_n-T_i}^{(i)}}\right)\right)^{M_{t_n}}\right).
\end{aligned}$$

The Laplace transform of a geometric random variable $Y$ of parameter $p$ is $\mathbb{E}\left(e^{-uY}\right) = \frac{p}{e^u-(1-p)}$, thus

$$\begin{aligned}
\mathcal{L}_{Z_{10}^{(n)}/n^q}(u) &= \mathbb{E}_{M_{t_n}}\left(\mathbb{E}_{T_i|M_{t_n}}\left(\frac{p_{10}(t_n-T_i)}{e^{\frac{u}{n^q}}-1+p_{10}(t_n-T_i)}\right)^{M_{t_n}}\right) \\
&= \mathbb{E}_{M_{t_n}}\left(\left(\int_0^{t_n}\frac{p_{10}(t_n-t)}{e^{\frac{u}{n^q}}-1+p_{10}(t_n-t)} * \frac{\lambda_{00}\mu_{\text{high}}N_t s_{10}(t_n-t)}{\Lambda(t_n)}\,\mathrm{d}t\right)^{M_{t_n}}\right) \\
&= \sum_{m=0}^{\infty}\left(\int_0^{t_n}\frac{p_{10}(t_n-t)}{e^{\frac{u}{n^q}}-1+p_{10}(t_n-t)} * \frac{\lambda_{00}\mu_{\text{high}}N_t s_{10}(t_n-t)}{\Lambda(t_n)}\,\mathrm{d}t\right)^m\frac{\Lambda(t_n)^m}{m!}e^{-\Lambda(t_n)} \\
&= e^{-\Lambda(t_n)}\sum_{m=0}^{\infty}\left(\int_0^{t_n}\frac{p_{10}(t_n-t)}{e^{\frac{u}{n^q}}-1+p_{10}(t_n-t)}\lambda_{00}\mu_{\text{high}}N_t s_{10}(t_n-t)\,\mathrm{d}t\right)^m\frac{1}{m!} \\
&= \exp\left(-\Lambda(t_n)\right)\exp\left(\int_0^{t_n}\lambda_{00}\mu_{\text{high}}N_t s_{10}(t_n-t)\frac{p_{10}(t_n-t)}{e^{\frac{u}{n^q}}-1+p_{10}(t_n-t)}\,\mathrm{d}t\right).
\end{aligned}$$

Then recalling that $\Lambda(t_n) = \int_0^{t_n}\lambda_{00}\mu_{\text{high}}N_t s_{10}(t_n-t)\,\mathrm{d}t$, we have

$$\mathcal{L}_{Z_{10}^{(n)}/n^q}(u) = \exp\left(\int_0^{t_n}\lambda_{00}\mu_{\text{high}}N_t s_{10}(t_n-t)\left(\frac{p_{10}(t_n-t)}{e^{\frac{u}{n^q}}-1+p_{10}(t_n-t)}-1\right)\mathrm{d}t\right)$$

and by replacing $p_{10}(t_n-t)$ and $s_{10}(t_n-t)$ by their expression we obtain:

$$\mathcal{L}_{Z_{10}^{(n)}/n^q}(u) = \exp\left(\int_0^{t_n}\lambda_{00}n^{\beta-1}e^{r_{00}t}r_{10}\frac{1-e^{\frac{u}{n^q}}}{e^{r_{10}(t-t_n)}(\lambda_{10}(1-\mu_{\text{low}})-d_{10}e^{\frac{u}{n^q}})-\lambda_{10}(1-\mu_{\text{low}})(1-e^{\frac{u}{n^q}})}\,\mathrm{d}t\right).$$

Using the change of variables $y = \frac{e^{r_{00}t}-1}{e^{r_{00}t_n}-1}$, $\mathrm{d}y = \frac{r_{00}e^{r_{00}t}}{e^{r_{00}t_n}-1}\,\mathrm{d}t$ leads to:

$$\begin{aligned}
\mathcal{L}_{Z_{10}^{(n)}/n^q}(u) &= \exp\left(-\frac{\lambda_{00}r_{10}}{r_{00}}n^{\beta-1}(n^{\alpha-\beta}-1)\frac{(e^{\frac{u}{n^q}}-1)}{\lambda_{10}(1-\mu_{\text{low}})-d_{10}e^{\frac{u}{n^q}}}\int_0^1\frac{\mathrm{d}y}{\frac{((n^{\alpha-\beta}-1)y+1)^{\frac{r_{10}}{r_{00}}}}{n^{\frac{r_{10}}{r_{00}}(\alpha-\beta)}}+\frac{\lambda_{10}(1-\mu_{\text{low}})(e^{\frac{u}{n^q}}-1)}{\lambda_{10}(1-\mu_{\text{low}})-d_{10}e^{\frac{u}{n^q}}}}\right) \\
&= \exp(g(n)I_n)
\end{aligned}$$

with

$$\begin{aligned}
g(n) &= -\frac{\lambda_{00}r_{10}}{r_{00}}n^{\beta-1}(n^{\alpha-\beta}-1)\frac{(e^{\frac{u}{n^q}}-1)}{\lambda_{10}(1-\mu_{\text{low}})-d_{10}e^{\frac{u}{n^q}}} \\
&\underset{n\to+\infty}{\sim}\frac{-u\lambda_{00}}{r_{00}}n^{\alpha-1-q}
\end{aligned}$$

and

$$I_n = \int_0^1 \frac{\mathrm{d}y}{\frac{((n^{\alpha-\beta}-1)y+1)^{\frac{r_{10}}{r_{00}}}}{n^{\frac{r_{10}}{r_{00}}(\alpha-\beta)}} + \frac{\lambda_{10}(1-\mu_{\text{low}})(e^{\frac{u}{n^q}}-1)}{\lambda_{10}(1-\mu_{\text{low}})-d_{10}e^{\frac{u}{n^q}}}}.$$

For $n \to \infty$, $I_n$ is equivalent to

$$I_n^* = \int_0^1 \frac{\mathrm{d}y}{y^c + a_n} = \int_0^\infty \frac{e^{-x}}{e^{-cx} + a_n}\,\mathrm{d}x$$

where $c = \frac{r_{10}}{r_{00}}$ and $a_n = \frac{\lambda_{10}(1-\mu_{\text{low}})u}{r_{10}n^q}$. We can compute this integral using the residue theorem [7]: we integrate the function $h(x) = \frac{e^{-x}}{e^{-cx}+a_n}$ on a rectangle with width $R$ and height $\frac{2\pi}{c}$ with bottom left corner on the origin. If we respectively name $I_R$, $K_R$, $J_R$ and $L_R$ the integrals along the four sides of the rectangle, then we have $I_R \underset{R\to\infty}{\to} I_n^*$, $J_R = -e^{-\frac{2i\pi}{c}} I_R$ and $J_R + K_R \underset{R\to\infty}{\to} 0$. As this rectangle contains one simple pole of $h(x)$ (for $x = -\frac{\ln(a_n)}{c} + \frac{i\pi}{c}$), we obtain:

$$I_n^*(1 - e^{-\frac{2i\pi}{c}}) = 2i\pi \mathrm{Res}\left(\frac{e^{-x}}{e^{-cx} + a_n}, -\frac{\ln(a_n)}{c} + \frac{i\pi}{c}\right)$$

$$I_n^* = \frac{2i\pi}{1 - e^{-\frac{2i\pi}{c}}} * \frac{e^{\frac{\ln(a_n)}{c} - \frac{i\pi}{c}}}{-ce^{-c(-\frac{\ln(a_n)}{c} + \frac{i\pi}{c})}}$$

$$I_n^* = \frac{2i\pi}{e^{\frac{i\pi}{c}} - e^{-\frac{i\pi}{c}}} * \frac{a_n^{\frac{1}{c}}}{ca_n}$$

$$I_n^* = \frac{\pi}{c\sin(\frac{\pi}{c})} a_n^{\frac{1}{c}-1}$$

It follows that

$$I_n \underset{n\to+\infty}{\sim} \frac{r_{00}\pi}{r_{10}\sin\left(\frac{r_{00}\pi}{r_{10}}\right)} \left(\frac{\lambda_{10}u}{r_{10}n^q}\right)^{\frac{r_{00}}{r_{10}}-1}$$

and

$$g(n)I_n \underset{n\to\infty}{\sim} \frac{-\lambda_{00}\pi}{\lambda_{10}\sin\left(\frac{r_{00}\pi}{r_{10}}\right)} \left(\frac{\lambda_{10}u}{r_{10}}\right)^{\frac{r_{00}}{r_{10}}} n^{\alpha-1-\frac{r_{00}}{r_{10}}q}.$$

Thus $\mathcal{L}_{Z_{10}^{(n)}/n^q}(u)$ has a finite limit when $n \to \infty$ for $q = (\alpha-1)\frac{r_{10}}{r_{00}}$ :

$$\mathcal{L}_{Z_{10}^{(n)}/n^{(\alpha-1)\frac{r_{10}}{r_{00}}}}(u) \underset{n\to\infty}{\to} \exp\left(-\frac{\lambda_{00}\pi}{\lambda_{10}\sin\left(\frac{r_{00}\pi}{r_{10}}\right)}\left(\frac{\lambda_{10}u}{r_{10}}\right)^{\frac{r_{00}}{r_{10}}}\right).$$

It follows that

$$Z_{10}^{(n)} \underset{n\to+\infty}{\sim} n^{(\alpha-1)\frac{r_{10}}{r_{00}}} Z_{10}$$

with $Z_{10}$ a random variable independent of $n$ with Laplace transform:

$$\mathcal{L}_{Z_{10}}(u) = \exp\left(-\frac{\lambda_{00}\pi}{\lambda_{10}\sin\left(\frac{r_{00}\pi}{r_{10}}\right)}\left(\frac{\lambda_{10}u}{r_{10}}\right)^{\frac{r_{00}}{r_{10}}}\right).$$

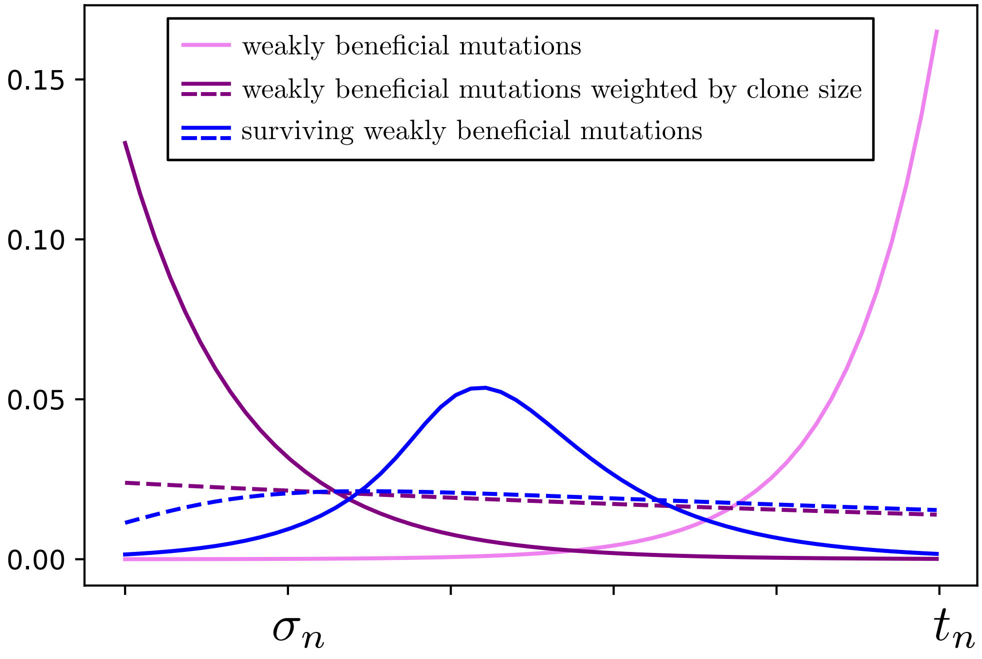

Figure 1: Distribution in time of weakly beneficial mutational origins during first cycle. Pink curve: total weakly beneficial mutation rate, purple curves: total weakly beneficial mutation rate weighted by final expected clone size, and blue curves: rate of emergence of weakly beneficial mutations that survive the first bottleneck. Solid curves are plotted for $r_{10} = 0.3$, dashed ones for $r_{10} = 0.18$ ($r_{00} = 0.17$, $n = 10^8$, $\alpha = 1.4$, $\beta = 0.9$).

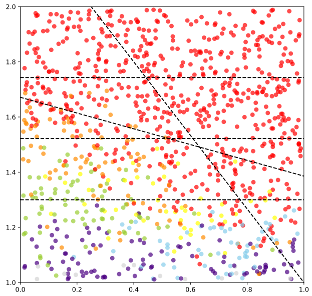

(a) Exponential growth, $n = 10^6$

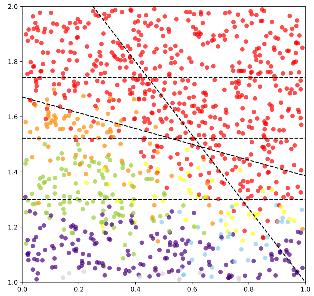

(b) Exponential growth, $n = 10^9$

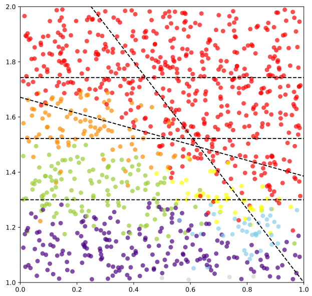

(c) Exponential growth, $n = 10^{12}$

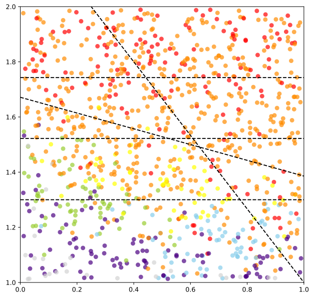

(d) Density-dependent growth, $n = 10^6$

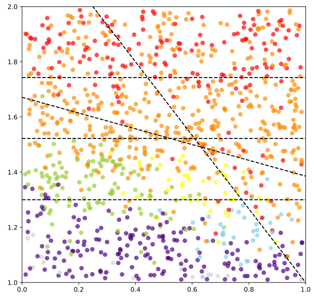

(e) Density-dependent growth, $n = 10^9$

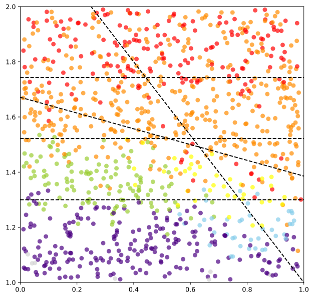

(f) Density-dependent growth, $n = 10^{12}$

Figure 2: Observed paths in simulations with exponential (a,b,c) or density-dependent (d,e,f) growth: each point corresponds to a simulation and is colored according to the observed scenario. Simulations were run for 7 cycles with $n = 10^6$ (a,d), $n = 10^9$ (b,e) or $n = 10^{12}$ (c,f). $\gamma = 2, \delta = 1.3, r_{00} = 0.2, r_{10} = 0.35, r_{01} = 0.9, r_{11} = 1$ and $d_{00} = d_{10} = d_{01} = d_{11} = 0.1$.

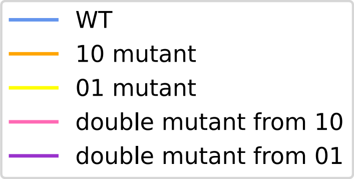

(a) Legend

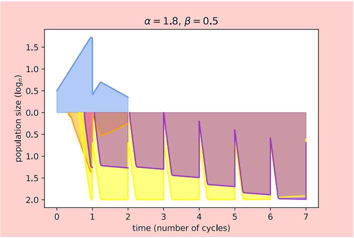

(b) Adaptation mainly from 01

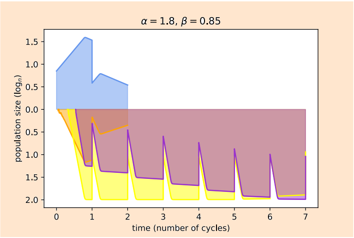

(c) Adaptation from 01

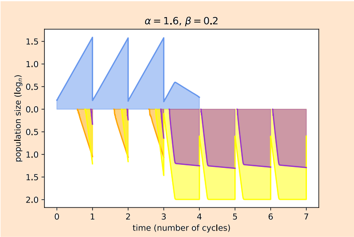

(d) Adaptation from 01

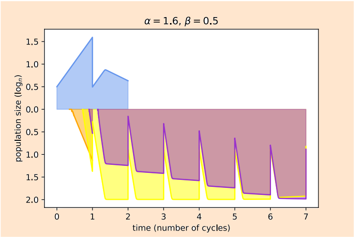

(e) Adaptation from 01

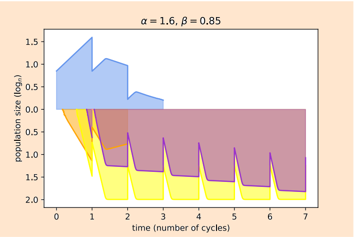

(f) Adaptation from 01

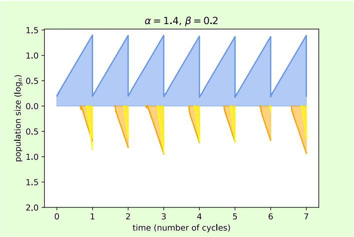

(g) No adaptation

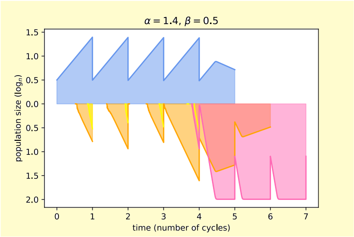

(h) Adaptation from 10

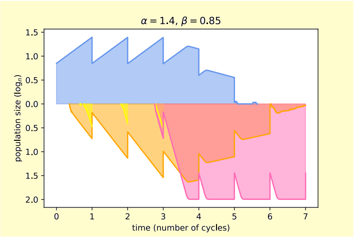

(i) Adaptation from 10

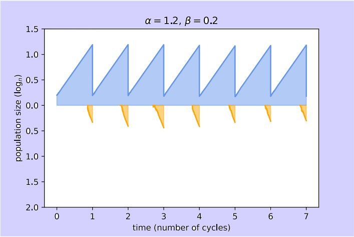

(j) No adaptation

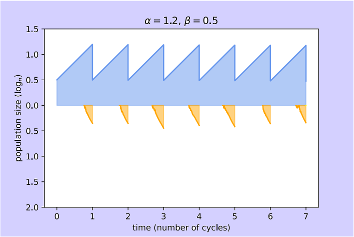

(k) No adaptation

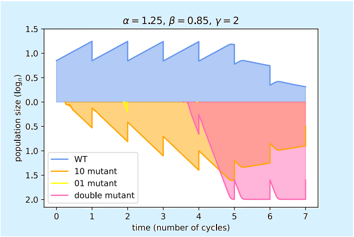

(l) Adaptation from 10

Figure 3: Examples of simulated trajectories for $(\alpha, \beta) \in \{1.25, 1.4, 1.6, 1.8\} \times \{0.2, 0.5, 0.85\}$, with density-dependence. WT population size is above, mutant population sizes are plotted below and superimposed. Each simulation was run for 7 cycles, with parameters $n = 10^9$, $\gamma = 2$, $\delta = 1.3$, $r_{00} = 0.2$, $r_{10} = 0.35$, $r_{01} = 0.9$, $r_{11} = 1$ and $d_{00} = d_{10} = d_{01} = d_{11} = 0.1$.

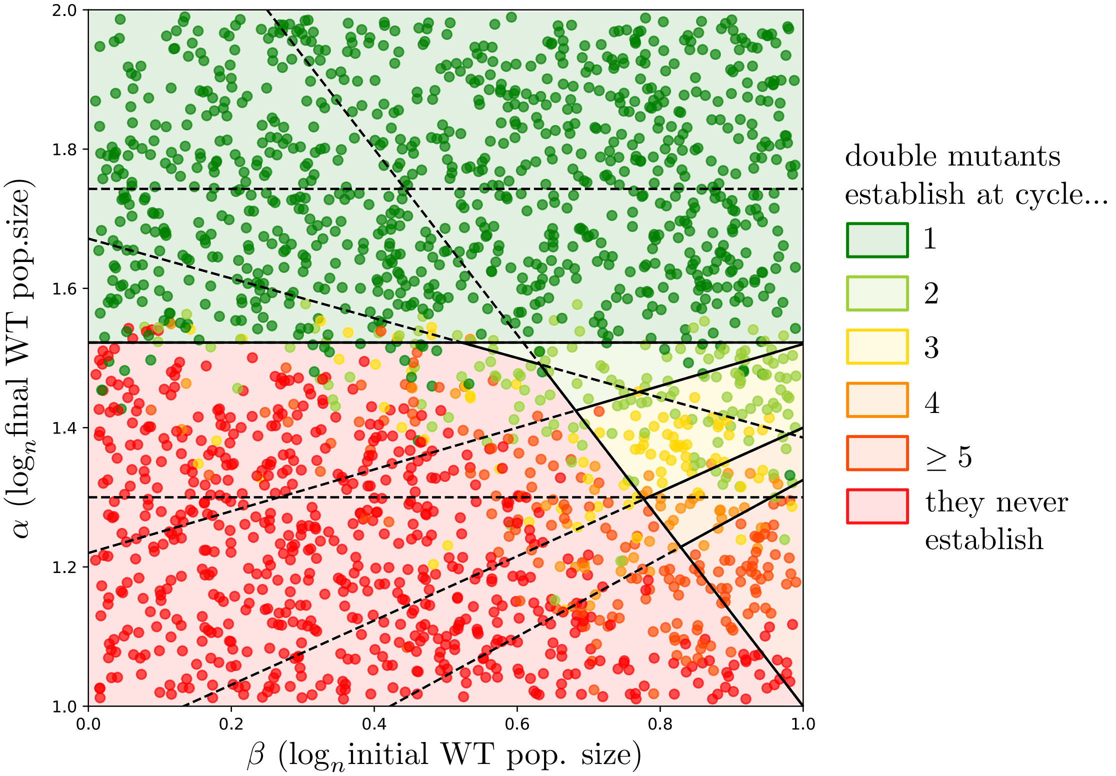

Figure 4: Observed cycles of double mutant emergence in 2,000 simulations with density-dependent growth. Simulations were run for 10 cycles with $n = 10^{12}$.
$\gamma = 2, \delta = 1.3, r_{00} = 0.2, r_{10} = 0.35, r_{01} = 0.9, r_{11} = 1$ and $d_{00} = d_{10} = d_{01} = d_{11} = 0.1$. Boundaries and background colors are theoretical predictions for exponential growth and large $n$.

[7] Lars Ahlfors. *Complex Analysis.* McGraw-Hill Inc., New York, 1979.
